# Robotic Antimicrobial Susceptibility Platform (RASP): A Next Generation Approach to One-Health Surveillance of Antimicrobial Resistance

**DOI:** 10.1101/2021.03.09.434587

**Authors:** Alec Truswell, Rebecca Abraham, Mark O’Dea, Zheng Zhou Lee, Terence Lee, Tanya Laird, John Blinco, Shai Kaplan, John Turnidge, Darren J. Trott, David Jordan, Sam Abraham

## Abstract

**Background:** Surveillance of antimicrobial resistance (AMR) is critical to reducing its wide-reaching impact. Its reliance on sample size invites solutions to longstanding constraints regarding scalability. A robotic platform (RASP) was developed for high-throughput AMR surveillance in accordance with internationally recognised standards (CLSI and ISO 20776-1:2019) and validated through a series of experiments.

**Methods:** Experiment A compared RASP’s ability to achieve consistent MICs to that of a human technician across eight replicates for four *E. coli* isolates. Experiment B assessed RASP’s agreement with human performed MICs across 91 *E. coli* isolates with a diverse range of AMR profiles. Additionally, to demonstrate its real-world applicability, the RASP workflow was then applied to five faecal samples where a minimum of 47 *E. coli* per animal (239 total) were evaluated using an AMR indexing framework.

**Results:** For each drug-rater-isolate combination in experiment A, there was a clear consensus of the MIC and deviation from the consensus remained within one doubling-dilution (the exception being gentamicin at two dilutions). Experiment B revealed a concordance correlation coefficient of 0.9670 (95%CI: 0.9670 - 0.9670) between the robot and human performed MICs. RASP’s application to the five faecal samples highlighted the intra-animal diversity of gut commensal *E. coli,* identifying between five and nine unique isolate AMR phenotypes per sample.

**Conclusions:** While adhering to internationally accepted guidelines, RASP was superior in throughput, cost and data resolution when compared to an experienced human technician. Integration of robotics platforms in the microbiology laboratory is a necessary advancement for future One-Health AMR endeavours.

## Introduction

Antimicrobial resistance (AMR) has been described as one of the greatest threats to global health and food security, with an estimated cumulative cost of 100 trillion USD by 2050.^1^ The major consequence of resistance is the escalating difficulty of successfully treating bacterial infections with the currently available limited array of antimicrobials. We now face the risk of returning to the pre-antibiotic era where bacterial infections in humans and animals were a major cause of mortality.^2^ Preserving the usefulness of existing antimicrobials by reducing selection and limiting the dissemination of resistant organisms is therefore a high priority. Consequently, a key component of management efforts is surveillance designed to keep authorities and clinicians aware of where and when resistance is present and evolving.^3^ Surveillance of AMR faces many challenges owing to the multi-host, multi-pathogen, multi-drug nature of the resistance phenomenon. Perhaps one of the most problematic aspects is the insidious nature of resistance - it emerges slowly without necessarily yielding expression of any outward signs and frequently has an impact at a place and time other than its origin. For surveillance to overcome these obstacles, careful attention needs to be paid to the choice of pairing of assays for measuring resistance in bacteria and the design of sample acquisition from animals, food and the environment.

Surveillance for AMR in animals and food is a well-established activity. The WHO has longstanding recommendations for the conduct of “integrated surveillance”, as part of the multifaceted-management for the control of resistance.^4^ Countries such as Denmark^5^ and the USA^6^ have been at the forefront of surveillance for AMR in animals and the food chain with programs running since as early as 1995. Since then, the basic approach in food-animals, which is now widely adopted throughout the developed world, has changed very little. A core component is the collection and analysis of data relating to the AMR profile of indicator bacteria (such as *Escherichia coli*) from various livestock and commodities.^7^ As a result of the high cost per isolate (of which laboratory processing contributes a significant portion), most national-level surveys typically only collect data on approximately 150-250 individually selected isolates per year from each livestock sector. While the resulting data provide a general overview on AMR in food-animals, the inferences that can be drawn are limited. In addition, recent data have demonstrated variation of *E. coli* resistance profiles within bacterial species from the same host,^8^ suggesting that reliable estimates of AMR require the collection of multiple isolates from a single host.

In order to effectively limit the emergence and spread of AMR within groupings of animals or humans, surveillance data must be strengthened to make it more relevant to antimicrobial stewardship.^9^ A paramount need is for the individuals responsible for antimicrobial stewardship in a setting to receive intelligence from surveillance describing the occurrence of resistance. To deliver such feedback, epidemiologically appropriate study design and sample size needs to be combined with cost-effective characterisation of much larger numbers of bacteria than are currently evaluated.^10^ At present the isolation of large numbers of bacterial colonies, subsequent AST with traditional techniques and genomic characterisation is not only cost-prohibitive but a severe drain on time and resources. Scaling up traditional methods is unattractive because it would inevitably introduce inaccuracy arising from fatigue of laboratory workers. However, the harnessing of high-throughput laboratory robotic platforms capable of handling individual colonies on an unprecedented scale and without the loss of accuracy from fatigue we associate with manual methods is a practical solution.

In this study, we describe the development of the next generation approach to surveillance for AMR using a Robotic Antimicrobial Susceptibility Platform (RASP) and demonstrate that the RASP method is elegantly and efficiently adapted to high-volume surveillance of commensal organisms. The system automates the process of bacterial isolation, identification (with or without reliance on spectrography), and AST for customised combinations of drugs. With this comes a much-needed improvement in the ability to design surveillance to meet objectives that are of practical relevance at the coalface of antimicrobial stewardship.

## Methods

### System overview

RASP was developed to assess phenotypic resistance in a manner comparable with major surveillance programs. Unlike other approaches to “scaling up” such as metagenomics, RASP keeps the nucleic acid of individual isolates available for study making it possible to understand if resistance is mediated by previously unidentified genes while avoiding biases due to the presence of extraneous nucleic acid from environmental organisms.

RASP is a customisable robotics platform that was designed, using available Tecan (Switzerland) and SciRobotics Ltd. (Israel) products, to be flexible and multifunctional, capable of bacterial isolation, isolate collection, preparation for identification and AST. A Tecan Freedom EVO® 150 base was combined with the SciRobotics carousel capable of holding both petri dishes and 96-well microplates. SciRobotics equipment was deployed on the robot deck, the PetriPlater™ to dispense samples and Pickolo™ software using image analysis to select colonies, in addition to a microplate absorbance reader for AST inoculum adjustment to deliver high-throughput surveillance (Figure S1). The robot has also been fully integrated with barcoding capabilities to track isolates from original source to phenotypic results and is integrable with any electronic laboratory information management system. The general workflow for this method is visualised in Figure 1 and the major steps detailed below. Homogenised samples are loaded onto the robot in a liquid format and either serially diluted or directly streaked onto agar. After incubation individual colonies are selected using Pickolo™, based on the explicit requirements of the user including colour, size and morphology. Individual colonies are transferred to a 96-well microplate and a deep-well plate in preparation for AST and isolate storage, respectively. MALDI-TOF identification can also be performed at this timepoint and as such, the robot includes a position for a MALDI-TOF target plate and is capable of both sample and matrix addition. AST is then performed using an entirely liquid based methodology adapted for the RASP platform from internationally established guidelines: Clinical Laboratory Standards Institute (CLSI)^11^ and ISO 20776-1:2019; including compliance to quality control criteria. AST drug panels, genotyping and whole genome sequencing can be prepared using the Freedom EVO genomics platform. While the above workflow was well suited to this current study, it should be noted that it represents one of many possibilities. RASP workflows are flexible and due to the modular design of hardware, can be adjusted to suit different bacterial species, sample types, culture media and drug panels.

**Figure 1.**
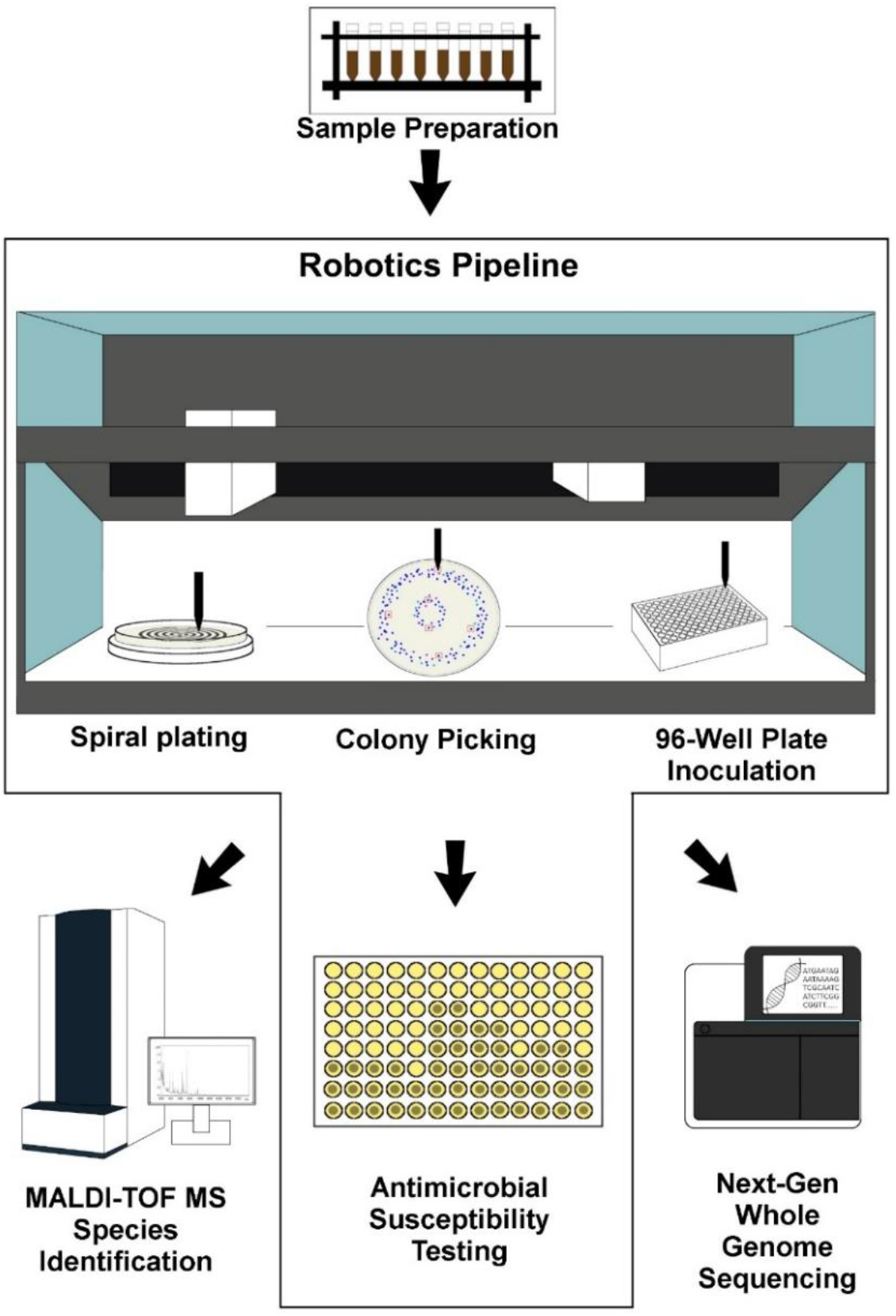
Overview of a typical RASP workflow. Any processes depicted outside the border occur externally to RASP.

### Bacterial Isolation

Two grams of faecal sample was homogenised in 18 mL PBS. Homogenised samples then underwent four 10-fold serial dilutions and was plated onto CHROMagar™ ECC (Edwards, MM1076) using the two-zone spiral plating protocol at 10^−3^ (outer zone) and 10^−4^ (inner zone) (Video S1). After overnight incubation at 37°C, Pickolo™ colony picking software was used to target and pick up to 48 presumptive *E. coli* colonies per plate based on their adherence to previously determined colour, size and circularity criteria (Video S2) and transferred to 96-well plates containing CAMHB (BD, 212322) for overnight growth at 37°C for storage and subsequent assays.

### Bacterial Identification

Isolates were prepared for MALDI-TOF identification by adding 40 μL of 90% Formic acid to the bacterial pellet from overnight culture. (Video S3). Species identification was then performed by MALDI-TOF adhering to manufacturer’s protocol (Bruker MALDI Biotyper Microflex LT/SH MALDI-MS running MBT Compass 4.1 Build 70 and flexControl 3.4 Build 135).

### Antimicrobial Susceptibility Testing

All isolates were grown from storage at −80°C onto sheep blood agar (Edwards, MM1120) overnight at 37°C. A second subculture was performed the following day, and the workflow diverged depending on the prospective rater.

#### RASP Platform

The robot protocol adjusted isolates from the overnight broth culture using an absorbance reader at 620 nm wavelength, to an absorbance equivalent to a McFarland standard (0.08 to 0.13). The McFarland standardised isolate was transferred to a deep well plate and diluted 1:20 in sterile water, and the drug plate was inoculated with 10 μL of isolate in 90 μL of CAMHB diluted drug (Video S4). For colony enumeration, the robot performed further dilutions (1:150 followed by another 1:150) on the previously diluted McFarland standard and plated 100 μL on sheep blood agar using a modification of the two-zone spiral dilution plating method; whereby the entire plate is inoculated as just one zone (lawn plating). Following an overnight incubation (16 – 20 hours at 37°C as per CLSI guidelines), drug plate results for both raters were read using the Sensititre Vizion plate reader system, and colony enumeration on sheep blood agar was performed and recorded using the Pickolo colony counting software.

#### Human Technician

Isolates intended for human performed susceptibility testing were subcultured on sheep blood agar, while isolates intended for robot performed susceptibility testing were subcultured in a flat-bottom 96-well plate (Nunc, 167008), each well filled with 220 μL (allowing for overnight evaporation of roughly 20 μL) CAMHB (BD, 212322). The human broth microdilution protocol followed CLSI guidelines, with the adjustment of isolates to a 0.5 McFarland standard using a nephelometer (Sensititre™) followed by a 1:100 dilution in CAMHB prior to inoculation of the drug plate using Sensititre’s AIM™ automated inoculation delivery system (50 μL inoculum into 50 μL of drug diluted in CAMHB). The colony enumeration quality control step was performed for each isolate on sheep blood agar as per CLSI guidelines with a 1:1000 dilution of inoculum in 0.9% saline and lawn plating on agar.

### Validation and Application of RASP

To evaluate the capacity of the robot and validate its use in AST, we applied the RASP robotic platform to the aforementioned workflow using a collection of *E. coli*; as *E. coli* is an ideal indicator organism and represents a ubiquitous component of surveillance for AMR in animals.^5^, ^6^, ^12^

### AST Validation Experiment A: Assessment of Repeatability

An experiment was conducted to validate the RASP platform’s AST protocol by comparing the ability of both human and robot rater to generate consistent minimum inhibitory concentrations (MICs) across replicates. Each rater performed AST on four *E. coli* isolates in octuplicate against the following antimicrobials: ampicillin, cefoxitin, ceftiofur, chloramphenicol, ciprofloxacin, colistin, ceftriaxone, florfenicol, gentamicin, streptomycin, trimethoprim/sulfamethoxazole and tetracycline. Isolates were derived from various sources including an American Type Culture Collection 25922, and one isolate of seagull^13^, porcine^14^ and cattle origin^15^, each with previously determined AMR profiles.

### AST Validation Experiment B: Assessment of Rater Agreeance

Experiment B was a breadth study whereby the RASP platform and an experienced human technician performed AST on a diverse range of isolates (n=91) with varying, previously characterised AMR profiles and host origins (seagull, porcine, cattle).^13–17^ The RASP platform was assessed based on its agreeance with the human technician in their determination of MICs for each isolate. Antimicrobials tested included: ampicillin, cefoxitin, ceftiofur, chloramphenicol, ciprofloxacin, colistin, ceftriaxone, florfenicol, gentamicin, streptomycin, trimethoprim/sulfamethoxazole and tetracycline.

### RASP Workflow Application: Antimicrobial Resistance Surveillance Index Scoring

To demonstrate the level of resolution achievable with RASP, five faecal samples were collected from pig pen floors (one sample per pen) and proceeded through the RASP workflow described above to yield AST data for 239 derivative *E. coli* isolates (minimum to 47 per sample) against six antimicrobials (ampicillin, ceftiofur, ciprofloxacin, gentamicin, tetracycline and trimethoprim sulfamethoxazole). A secondary classification and visualisation script devised in Stata^18^ was applied to the 239 isolates to generate a graphical representation of intra-animal and intra-herd diversity, as well as to assign an index score representative of the threat posed by their AMR burden. The indexing scheme assigned weights (w) to antimicrobials informed by national^19^ and international^20^ guidelines whereby the antimicrobials deemed more important to human health received a higher weighting. Antimicrobial weights were as follows: ampicillin (w=1), ceftiofur (w=3), ciprofloxacin (w=4), gentamicin (w= 2), tetracycline (w=1) and trimethoprim sulfamethoxazole (w=2). In this study the maximum possible AMR score was 13, indicating resistance to all tested antimicrobials.

### Throughput Comparison Analysis

An analytical component of this study sought to compare the time required by a human technician utilising current methods, versus a robot-assisted technician, to process 20 homogenised faecal samples (deriving 960 isolates) from isolation through to phenotypic characterisation. A simulated workflow was developed for each technician based on average times taken to perform each task within the workflow and applied to the above scenario. The human technician was allowed the use of ‘modern’ laboratory implements such as Sensititre’s auto-inoculator and Vizion plate reader systems, pre-ordered ready-to-use drug plates, and a MALDI-TOF, while the robot assisted technician had the two previously discussed Tecan robots, a MALDI-TOF and a Sensititre Vizion plate reader system at their disposal.

## Results

### AST Validation Experiment A: Assessment of Repeatability

In Experiment A, when comparing the human (H) and robot (R) MIC results for each isolate, the mode MIC of an isolate across both raters against a specific antimicrobial was labelled the ‘consensus MIC’ and any deviation from the consensus MIC deemed a departure from the truest result (Figure 2). Of the drug-rater-antimicrobial combinations tested (see Figure S2 for all combinations), the majority show all eight replicates having achieved the same MIC result, and furthermore demonstrate agreeance between the two raters; all isolates and replicates tested against tetracycline achieved the same MIC between both isolate and rater. Of the cases where deviation from the consensus MIC was seen, it was limited to one doubling-dilution (the exception being gentamicin at two dilutions) and there remained moderate agreement between the two raters, with similar distributions of replicates across MIC results.

**Figure 2.**
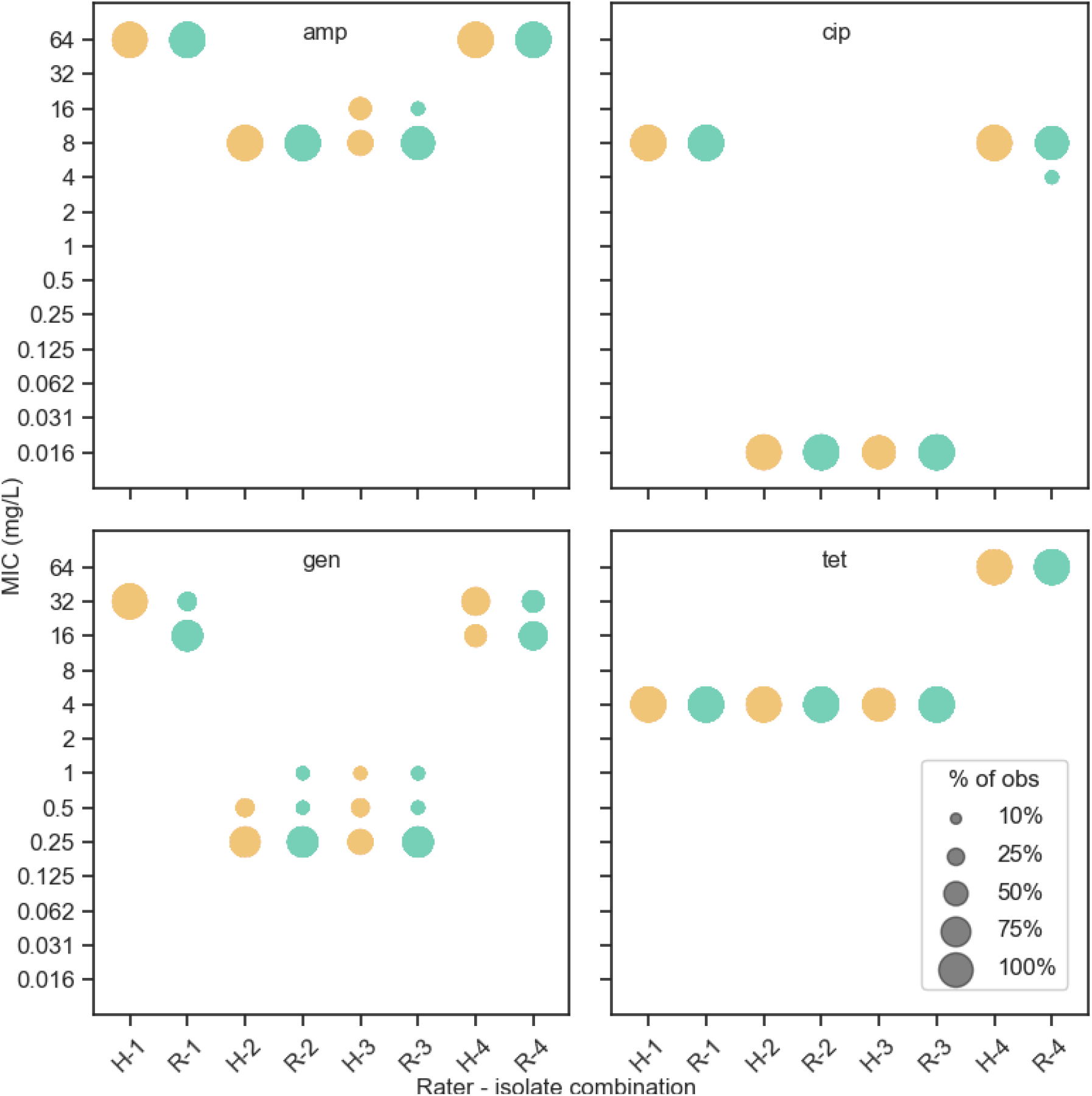
Experiment A: comparison of human (‘H’) and robot (‘R’) minimum inhibitory concentration (MIC) results of four fully susceptible *E. coli* isolates each replicated eight times per rater, tested against a panel of antimicrobials. Each isolate-rater designation displays the MIC results for all eight replicates of that combination. Antimicrobials: amp, ampicillin; cip, ciprofloxacin; gen, gentamicin; tet, tetracycline.

### AST Validation Experiment B: Assessment of Rater Agreement

In Experiment B, the majority of combinations of isolates and drugs (67.6 %) tested showed complete agreement between the two human and robot raters while 23.4% had a discrepancy in MIC result by one, which is acceptable according to international guidelines, and 9.0% had a discrepancy of two or more doubling dilutions (Figure 3). The greatest discrepancy in MIC between raters was a difference of nine dilutions between the two results and represented 0.26% of the isolate-drug combinations tested. The overall agreement for isolate-drug combinations between robot and human was summarised by a concordance correlation coefficient of 0.9670 (95%CI: 0.9670 - 0.9670), N: 1092.

**Figure 3.**
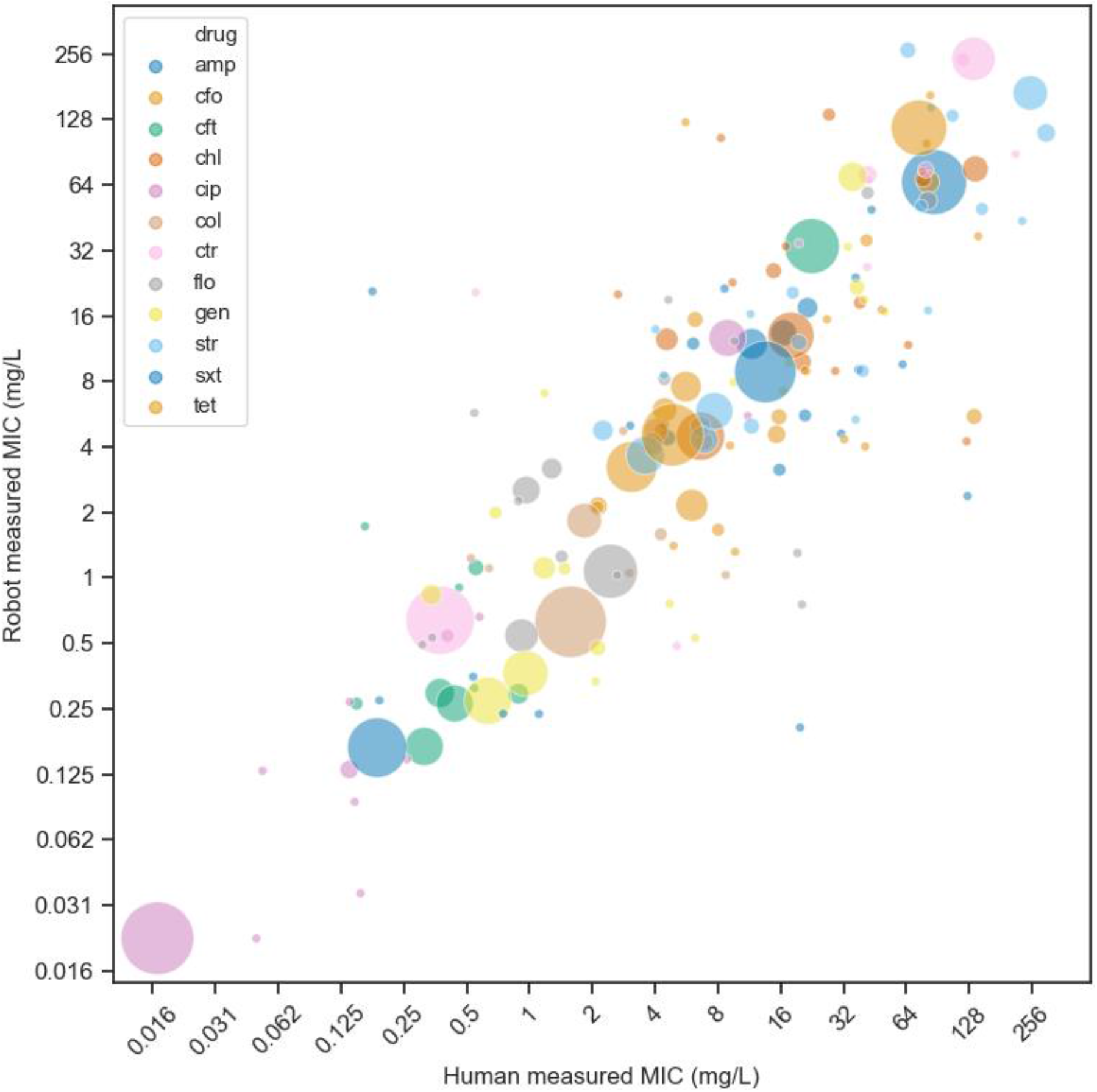
Experiment B: Paired minimum inhibitory concentration (MIC) results on 96 isolates with 12 drugs directly comparing human and robot measurements, marker size represents proportion of paired observations for the isolate-drug combination. Antimicrobials: amp, ampicillin; cfo, cefoxitin; cft, ceftiofur; chl, chloramphenicol; cip, ciprofloxacin; col, colistin; ctr, ceftriaxone; flo, florfenicol; gen, gentamicin; str, streptomycin; sxt, trimethoprim/sulfamethoxazole; tet, tetracycline.

### RASP Workflow Application: Antimicrobial Resistance Surveillance Index Scoring

All homogenised faecal samples that were plated by RASP on CHROMagar™ ECC selective agars yielded growth of at least 48 single colonies matching the expected blue appearance of *E. coli* at one or both dilutions. Isolates were successfully identified as *E. coli* using RASP’s MALDI-TOF preparation protocol, all with high identification scores; the lowest identification score achieved was 2.2 which qualified within Bruker’s highest confidence range.

Samples from all animals exhibited a diverse range of AMR profiles amongst isolates, including resistance to antimicrobials tested (Figure 4). All animals, except one, had isolates representing an AMR index of zero. The highest AMR index recorded was 13, comprising resistance to all tested antimicrobials. A relatively balanced distribution of AMR indices was seen for the 48 isolates from most animals; the exception being pig ‘E’, from which isolates were heavily skewed towards low AMR indices, with a high density achieving an AMR index of one.

**Figure 4.**
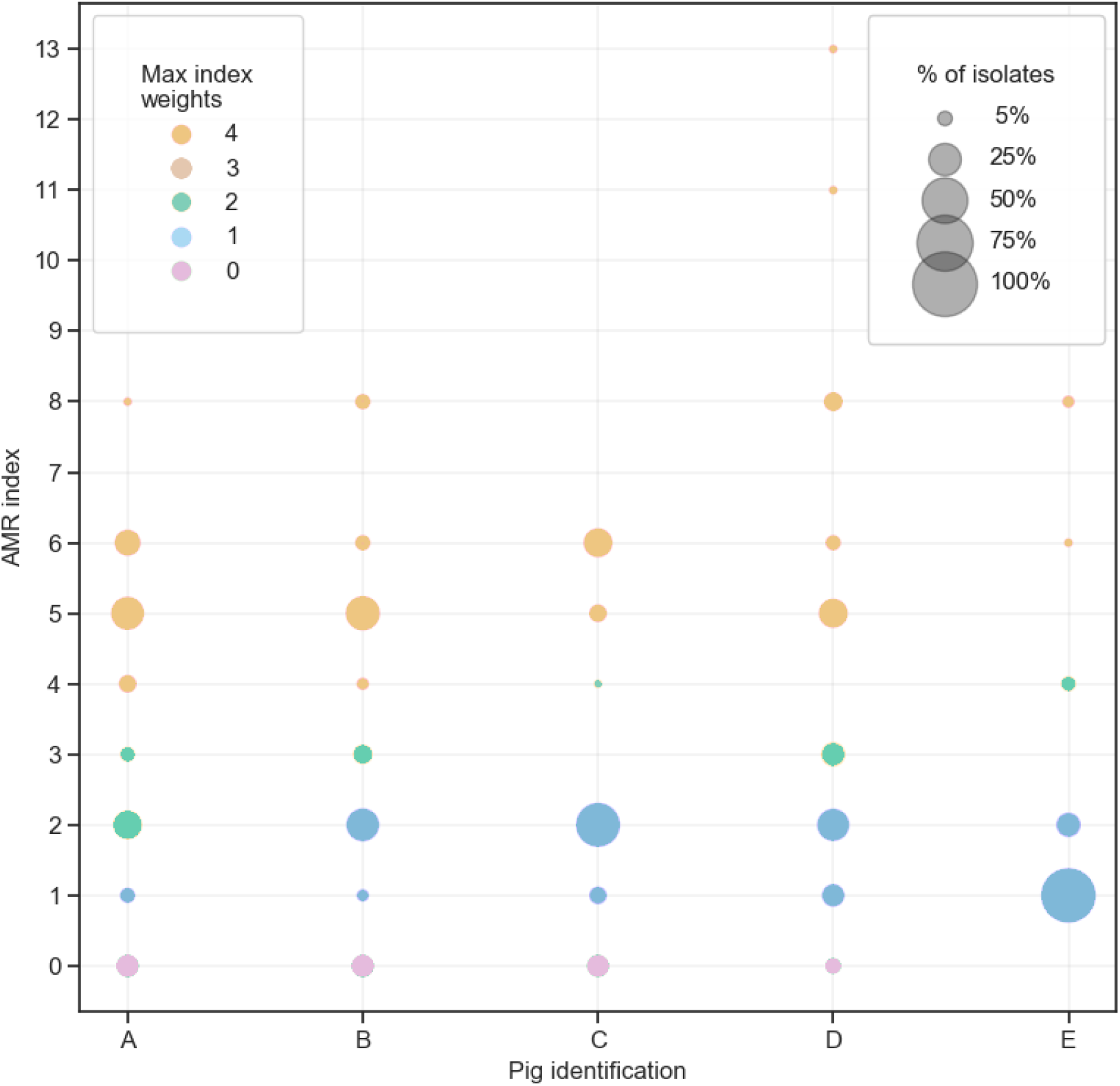
Antimicrobial resistance indexing scheme applied to 239 porcine *E. coli* isolates (48 from pigs A, B C and E, 47 from pig D) using the following antimicrobial risk-weightings (w): ampicillin (w=1), ceftiofur (w=3), ciprofloxacin (w=4), gentamicin (w= 2), tetracycline (w=1) and trimethoprim sulfamethoxazole (w=2). Colours are representative of the highest weighted resistance present for an isolate.

### Throughput Comparison Analysis

It was estimated that the human technician would take 30 days to process these samples compared to nine days for a robot-assisted technician (Figure S3). It’s important to state that the robots were not in continuous use during this workflow, meaning there was time where they could also have been utilised for other tasks.

## Discussion

The impetus to transition microbiology from a manual-labour-centric profession to one embracing automation is nothing novel. Automation offers the same advantage it has provided industry for decades; the augmentation of processes to increase efficiency and throughput; and this statement holds true for the application of the RASP platform to a conventional microbiology workflow. Typically, several technicians would be required in the manual workflow depicted in Figure S3 to prevent the occurrence of issues such as fatigue, and even still it is unlikely that optimal pace and quality of work would be maintained for the entirety of the workflow; a problem to which the robot is immune. The utilization of robotics instead allows staff to be diverted from monotonous and error-prone tasks to more cognitively intensive ones such as data analysis and project management.^21^, ^22^ These increases to throughput are only significant, however, if the solution is financially viable. It is therefore important to note that the improvement in processing efficiency is just one of several ways by which automation alleviates sample processing costs; savings are also seen on materials and reagents as a result of the transition to liquid-based methodologies (90% estimated cost reduction). In consideration of the initial financial outlay of purchasing the RASP platform (400,000 AUD; equivalent to approximately 300,000 USD or 260,000 EUR), we determined that approximately 9,000 samples would need to be processed for the above savings to equate to this initial cost. The incorporation of robotics into the routine microbiological assays described in this study is the technological leap required to elevate the standard of future One-Health AMR surveillance.

While currently available laboratory robotic systems (e.g. VITEK^®^, Microscan, WASPLab^®^ and BD Phoenix™) do offer the automation of isolation, identification or antimicrobial susceptibility testing, in comparison to RASP they utilise largely inflexible (in terms of pre-determined AST panels and characterisation of a single isolate)^23^ procedures to generate this information. We have developed the first integrated system – from isolation to AMR profile production that meets international guidelines and demonstrated that the method meets or exceeds the standards required. The Freedom EVO microbiology robot and by association the RASP protocol described here, utilises broth microdilution testing and MALDI-TOF integration (both gold-standard microbiological procedures) to capitalise on the higher throughput and resolution of data offered by its liquid-adapted methods.

AST completed by the robot was comparable to human generated data with similar patterns of MIC variance observed. AST in its current format is highly variable as can be seen by the wide range of MIC values acceptable for highly tested ATCC control strains.^11^ This phenomenon was well depicted in Experiment A, where under well-controlled, faithfully replicated conditions, MIC results from replicates of the same isolate spanning two or three doubling dilutions were commonly observed. The symmetry of MIC distributions between raters in these instances suggests that the source of this variation is biological in nature, due perhaps, to phenomena such as isolates expelling their plasmid, delayed expression of AMR genes or natural assay variations, as opposed to a technical failure by either rater.

Due to the restrictions of conventional methods considered above, most surveys will collect information for approximately 200 isolates per year per country, which limits the inferences that can be drawn. For example, with respect to the ability to detect early-emergence of resistance to a key drug, a simple application of binomial probability (expressed as: P(X>0) | X~Bin(200, 0.01)) demonstrates that a survey relying on 200 isolates has a probability of only 13.4% of detecting any positives if only 1% of isolates in the population have the resistance trait of interest. This calculation substantially underestimates the true number of isolates required during surveillance because reliance on a multi-staged sampling design (selection of herds, then animals, then isolates) is essential and carries a cost in accuracy created by the design effect.^24^ The AMR indexing experiment included in this study supports other data^8^, ^25^ in confirming that variation of *E. coli* resistance profiles within bacterial species from the same host and same group of animals is commonplace. The high level of resolution available from RASP for describing resistance in microbial populations, and its ability to yield an affordable increase in the number of samples and isolates assessed, was demonstrated in the final validation experiment in this study by the diversity of AMR profiles. In this final study only 0.4% (n=1) of tested isolates had an AMR index of 13, 1.7% (n=4) had resistance to gentamicin and 1.3% (n=3) had resistance to ceftiofur. There was also a corresponding variation in the number of AMR profiles for *E. coli* within a host: there being from 6-9 distinct variants detected in individuals, and more would likely be detected with higher sample sizes. With the availability of RASP there is an unprecedented opportunity to strengthen the management of AMR by shifting from a focus of interpretation adapted to clinical isolates to one that expressly accommodates the diversity of resistance in hosts at every level of organisation. Data generated on an expanded scale by RASP therefore presents an opportunity to provide more meaningful guidance on antimicrobial stewardship at the farm level through to national and international policy.

## Conclusion

RASP and technologies alike, unlock a higher calibre of AMR surveillance by overcoming longstanding constraints to scalability. The flexibility of RASP permits its application well beyond the scope demonstrated here, to the diverse range of bacterial landscapes encountered in the One-Health system. This study saw RASP’s workflow benchmarked against that of a contemporary laboratory, where it was demonstrated to be equivalent to an experienced human technician but proved superior in throughput, endurance and cost. It is critical that microbiology harnesses robotic platforms like RASP if we are to resist the current trajectory of AMR.

## Author Contributions

A. T., R. A., Z. L., T. L. and S. A. designed and performed the plating and colony picking experiments. A. T., R. A., J. B. and S. A. designed and performed the antimicrobial susceptibility testing experiments. J. T. and D. T. provided consultation on the antimicrobial susceptibility testing experiments as well as assisting with its validation according to international guidelines. S. K. developed the RASP platform adapting the hardware and software for microbiological use and provided consultation throughout the study. S. A., D. J., R. A., M. O. and D. T. conceived the project and S. A., R. A., M. O. and D. J. supervised the project. A. T., R. A., D. J., T. Lee. and S. A. performed data analysis. A. T., R. A., D. J. and S. A. wrote the initial draft of the manuscript, T. L., A. T., R. A. and S. A. designed figures and all authors contributed to editing and proof reading of the manuscript.

## Transparency Declaration

The authors declare no competing interests.

