## Supplementary Figures S1 - S3 for "Robotic Antimicrobial Susceptibility Platform (RASP): A Next Generation Approach to One-Health Surveillance of Antimicrobial Resistance"

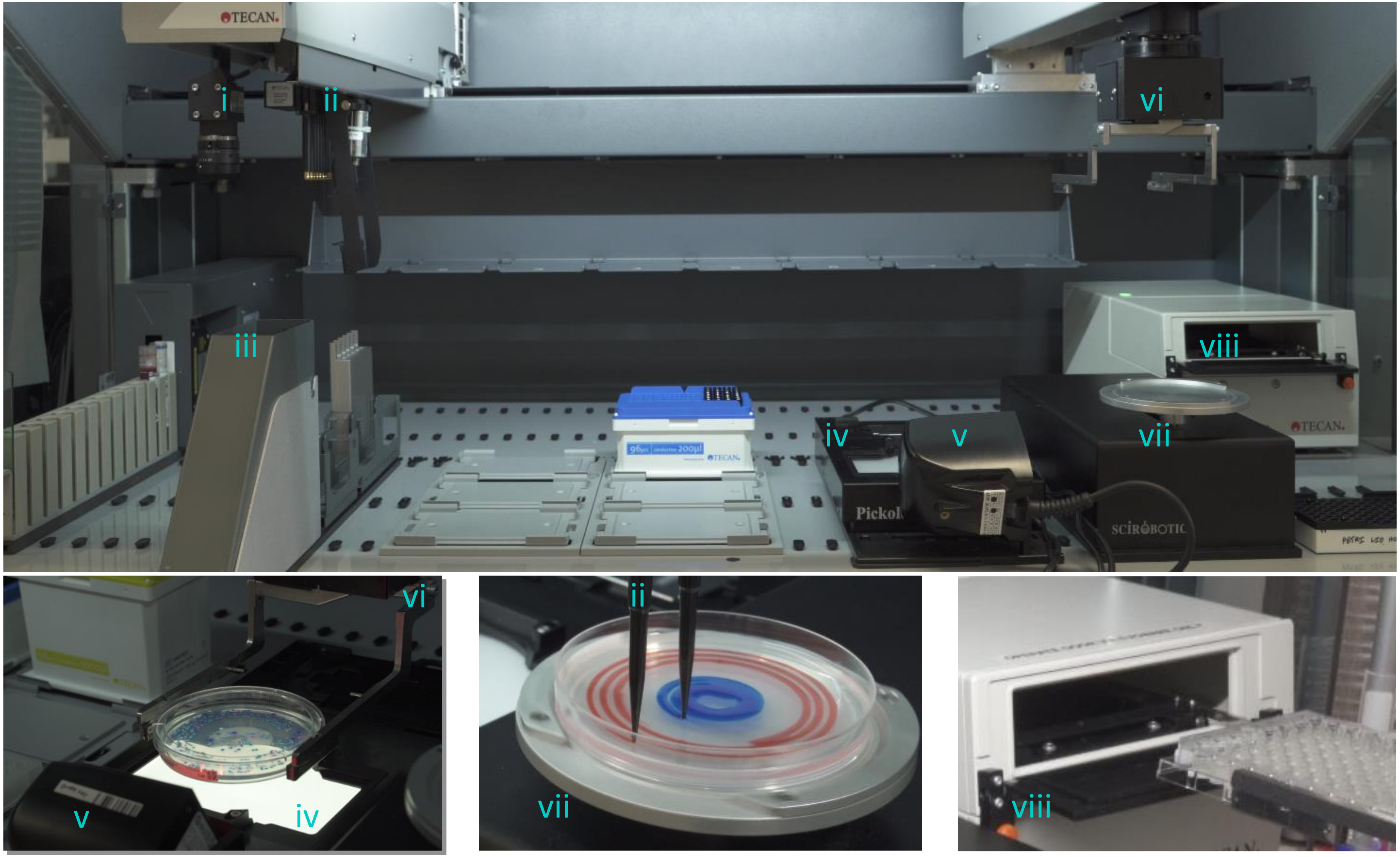


**Figure S1** | (a) Robotic Antimicrobial Susceptibility Platform (RASP) layout: i) camera, ii) liquid handling arm used for pipetting and colony picking, iii) waste chute, iv) lighting platform for plate imaging, v) barcode scanner, vi) plate handling arm, vii) agar inoculation platform, viii) spectrophotometer.

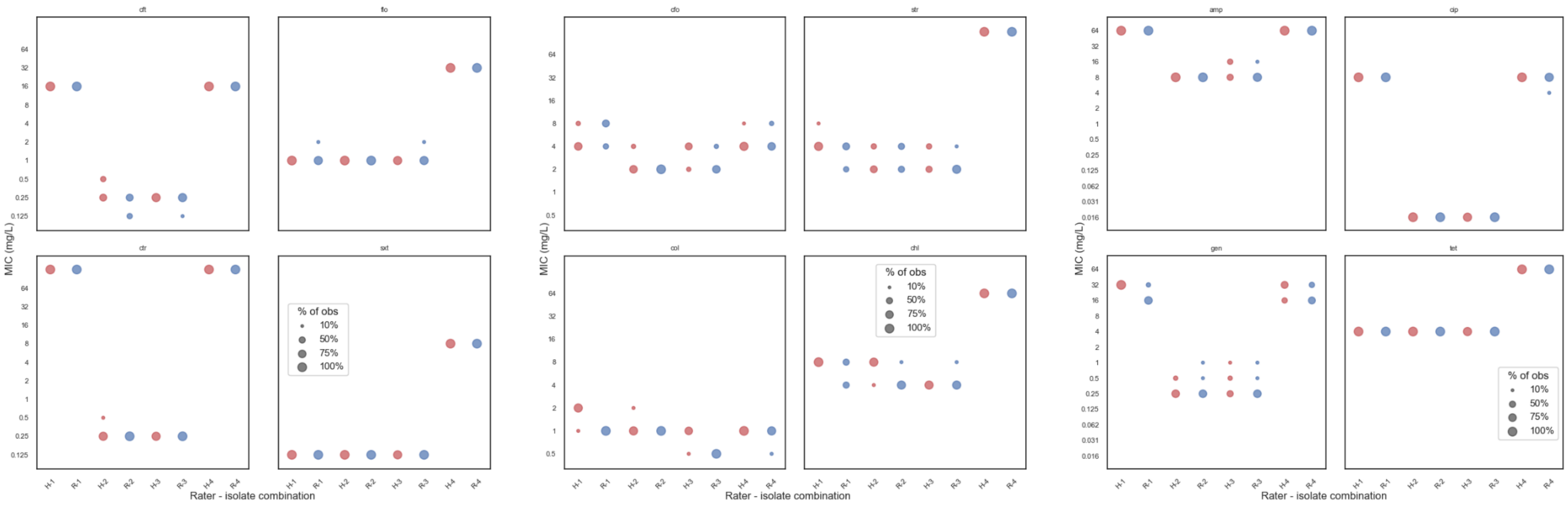


**Figure S2** | Experiment A: comparison of human ('H') and robot ('R') minimum inhibitory concentration (MIC) results of four fully susceptible *E. coli* isolates each replicated eight times per rater, tested against a panel of antimicrobials. Each isolate-rater designation displays the MIC results for all eight replicates of that combination. Antimicrobials: amp, ampicillin; cfo, cefoxitin; cft, ceftiofur; chl, chloramphenicol; cip, ciprofloxacin; col, colistin; ctr, ceftriaxone; flo, florfenicol; gen, gentamicin; str, streptomycin; sxt, trimethoprim/sulfamthoxazole; tet, tetracycline.

| Day One | | Hour One | Hour Two | Hour Three | Hour Four | Hour Five | Hour Six | our Seven | our Eight |
|---|---|---|---|---|---|---|---|---|---|
| | | 10 20 30 40 50 60 | 10 20 30 40 50 60 | 10 20 30 40 50 60 | 10 20 30 40 50 60 | 10 20 30 40 50 60 | 10 20 30 40 50 60 | 10 20 30 40 50 60 | 10 20 30 40 50 60 |
| Conventional Laboratory | Human (Sensititre Assisted) | Dilution Plating: Agar | | | | | | | |
| RASP Laboratory | Robot (Micro) | Dilution Plating: Agar | | | | | | | |
| | Robot (Prep) | | | | | | | | |
| | Human (Sensititre Assisted) | | | | | | | | |

| Day Two | | Hour One | Hour Two | Hour Three | Hour Four | our Five | ur Six | r Seven | Eight |
|---|---|---|---|---|---|---|---|---|---|
| | | 10 20 30 40 50 60 | 10 20 30 40 50 60 | 10 20 30 40 50 60 | 10 20 30 40 50 60 | 10 20 30 40 50 60 | 10 20 30 40 50 60 | 10 20 30 40 -0 0 | 10 |
| Conventional Laboratory | Human (Sensititre Assisted) | Colony Picking and First Subculture: Agar | | | | | | | |
| RASP Laboratory | Robot (Micro) | Colony Picking and Subculture | | | | | | culture (1-160) erve | |
| | Robot (Prep) | | | | | | | | |
| | Human (Sensititre Assisted) | | | | | | | | |

| Day Three | | Hour One | Hour Two | Hour Three | Hour Four | our Five | ur Six | r Seven | Eight |
|---|---|---|---|---|---|---|---|---|---|
| | | 10 20 30 40 50 60 | 10 20 30 40 50 60 | 10 20 30 40 50 60 | 10 20 30 40 50 60 | 10 20 30 40 50 60 | 10 20 30 40 50 60 | 10 !0 !0 0 0 ) | 10 |
| Conventional Laboratory | Human (Sensititre Assisted) | Colony Picking and First Subculture: Agar | MALDI-TOF Plate Preparation | | | | | | |
| [illegible] | Robot (Prep) | Master Drug Plate MIC Plate Preparation | | | | | | | |
| | Human (Sensititre Assisted) | | | | | | | | ilture (161-320) |

| Day Four | | Hour One | Hour Two | Hour Three | Hour Four | our Five | ur Six | r Seven | Eight |
|---|---|---|---|---|---|---|---|---|---|
| | | 10 20 30 40 50 60 | 10 20 30 40 50 60 | 10 20 30 40 50 60 | 10 20 30 40 50 60 | 10 20 30 40 50 60 | 10 20 30 40 50 60 | 10 !0 !0 0 0 ) | 10 |
| Conventional Laboratory | Human (Sensititre Assisted) | MALDI-TOF Plate Preparation | | | | | eserve | | |
| | Robot (Prep) | Master Drug Plate MIC Plate Preparation | | | | | | | |
| | Human (Sensititre Assisted) | Susceptibility Testing | | | | | | | [illegible] |

| Day Five | | Hour One | Hour Two | Hour Three | Hour Four | Hour Five | Hour Six | Hour Seven | Hour Eight |
|---|---|---|---|---|---|---|---|---|---|
| | | 10 20 30 40 50 60 | 10 20 30 40 50 60 | 10 20 30 40 50 60 | 10 20 30 40 50 60 | 10 20 30 40 50 60 | 10 20 30 40 50 60 | 10 20 30 40 50 60 | 10 20 30 40 50 60 |
| Conventional Laboratory | Human (Sensititre Assisted) | Preserve | | | | | | | |
| RASP Laboratory | Robot (Micro) | | | | | | | | |
| | Robot (Prep) | Master Drug Plate MIC Plate Preparation | | | | | | | |
| | Human (Sensititre Assisted) | Susceptibility Testing | | | | | | | [illegible] |

| Day Six | | Hour One | Hour Two | Hour Three | Hour Four | Hour Five | Hour Six | Hour Seven | Hour Eight |
|---|---|---|---|---|---|---|---|---|---|
| | | 10 20 30 40 50 60 | 10 20 30 40 50 60 | 10 20 30 40 50 60 | 10 20 30 40 50 60 | 10 20 30 40 50 60 | 10 20 30 40 50 60 | 10 20 30 40 50 60 | 10 20 30 40 50 60 |
| Conventional Laboratory | Human (Sensititre Assisted) | Preserve | | | | | | | |
| RASP Laboratory | Robot (Micro) | | | | | | | | |
| | Robot (Prep) | Master Drug Plate MIC Plate Preparation | | | | | | | |
| | Human (Sensititre Assisted) | Susceptibility Testing | | | | | | | [illegible] |

| Day Seven | | Hour One | Hour Two | Hour Three | Hour Four | Hour Five | Hour Six | Hour Seven | Hour Eight |
|---|---|---|---|---|---|---|---|---|---|
| | | 10 20 30 40 50 60 | 10 20 30 40 50 60 | 10 20 30 40 50 60 | 10 20 30 40 50 60 | 10 20 30 40 50 60 | 10 20 30 40 50 60 | 10 20 30 40 50 60 | 10 20 30 40 50 60 |
| Conventional Laboratory | Human (Sensititre Assisted) | Preserve | | | | | | | |
| RASP Laboratory | Robot (Micro) | | | | | | | | |
| | Robot (Prep) | Master Drug Plate MIC Plate Preparation | | | | | | | |
| | Human (Sensititre Assisted) | Susceptibility Testing | | | | | | | [illegible] |

| Day Eight | | Hour One | Hour Two | Hour Three | Hour Four | our Five | ur Six | r Seven | Eight |
|---|---|---|---|---|---|---|---|---|---|
| | | 10 20 30 40 50 60 | 10 20 30 40 50 60 | 10 20 30 40 50 60 | 10 20 30 40 50 60 | 10 20 30 40 50 60 | 10 20 30 40 50 60 | 10 !0 !0 0 0 ) | 10 |
| Conventional Laboratory | Human (Sensititre Assisted) | Second Subculture (1-96): AST | | | | | | | |
| | Robot (Prep) | Master Drug Plate MIC Plate Preparation | | | | | | | |
| | Human (Sensititre Assisted) | Susceptibility Testing | | | | | | | |

| Day Nine | | Hour One | Hour Two | Hour Three | Hour Four | our Five | ur Six | r Seven | Eight |
|---|---|---|---|---|---|---|---|---|---|
| | | 10 20 30 40 50 60 | 10 20 30 40 50 60 | 10 20 30 40 50 60 | 10 20 30 40 50 60 | 10 20 30 40 50 60 | 10 20 30 40 50 60 | 10 !0 !0 0 0 ) | 10 |
| Conventional Laboratory | Human (Sensititre Assisted) | Antimicrobial Susceptibility Testing (1-96) | | | | | | | |
| RASP Laboratory | Robot (Micro) | | | | | | | | |
| | Robot (Prep) | | | | | | | | |
| | Human (Sensititre Assisted) | Susceptibility Testing | Result Reading [illegible] | | | | | | |

| Day Ten | | Hour One | Hour Two | Hour Three | Hour Four | our Five | ur Six | r Seven | Eight |
|---|---|---|---|---|---|---|---|---|---|
| | | 10 20 30 40 50 60 | 10 20 30 40 50 60 | 10 20 30 40 50 60 | 10 20 30 40 50 60 | 10 20 30 40 50 60 | 10 20 30 40 50 60 | 10 !0 !0 0 0 ) | 10 |
| Conventional Laboratory | Human (Sensititre Assisted) | Susceptibility Testing Result Reading (1-96) | | | | | | | nd Subculture (97-192): AST |
| RASP Laboratory | Robot (Micro) | | | | | | | | |
| | Robot (Prep) | | | | | | | | |
| | Human (Sensititre Assisted) | | | | | | | | |

| Day Eleven | | Hour One | Hour Two | Hour Three | Hour Four | our Five | ur Six | r Seven | Eight |
|---|---|---|---|---|---|---|---|---|---|
| | | 10 20 30 40 50 60 | 10 20 30 40 50 60 | 10 20 30 40 50 60 | 10 20 30 40 50 60 | 10 20 30 40 50 60 | 10 20 30 40 50 60 | 10 !0 !0 0 0 ) | 10 |
| Conventional Laboratory | Human (Sensititre Assisted) | Antimicrobial Susceptibility Testing (97-192) | | | | | | | |
| RASP Laboratory | Robot (Micro) | | | | | | | | |
| | Robot (Prep) | | | | | | | | |
| | Human (Sensititre Assisted) | | | | | | | | |

| Day Twelve | | Hour One | Hour Two | Hour Three | Hour Four | our Five | ur Six | r Seven | Eight |
|---|---|---|---|---|---|---|---|---|---|
| | | 10 20 30 40 50 60 | 10 20 30 40 50 60 | 10 20 30 40 50 60 | 10 20 30 40 50 60 | 10 20 30 40 50 60 | 10 20 30 40 50 60 | 10 !0 !0 0 0 ) | 10 |
| Conventional Laboratory | Human (Sensititre Assisted) | Susceptibility Testing Result Reading (97-192) | | | | | | | nd Subculture (193-288): AST |
| RASP Laboratory | Robot (Micro) | | | | | | | | |
| | Robot (Prep) | | | | | | | | |
| | Human (Sensititre Assisted) | | | | | | | | |

| Day Thirteen | | Hour One | Hour Two | Hour Three | Hour Four | our Five | ur Six | r Seven | Eight |
|---|---|---|---|---|---|---|---|---|---|
| | | 10 20 30 40 50 60 | 10 20 30 40 50 60 | 10 20 30 40 50 60 | 10 20 30 40 50 60 | 10 20 30 40 50 60 | 10 20 30 40 50 60 | 10 !0 !0 0 0 ) | 10 |
| Conventional Laboratory | Human (Sensititre Assisted) | Antimicrobial Susceptibility Testing (193-288) | | | | | | | |
| RASP Laboratory | Robot (Micro) | | | | | | | | |
| | Robot (Prep) | | | | | | | | |
| | Human (Sensititre Assisted) | | | | | | | | |

| Day Fourteen | | Hour One | Hour Two | Hour Three | Hour Four | our Five | ur Six | r Seven | Eight |
|---|---|---|---|---|---|---|---|---|---|
| | | 10 20 30 40 50 60 | 10 20 30 40 50 60 | 10 20 30 40 50 60 | 10 20 30 40 50 60 | 10 20 30 40 50 60 | 10 20 30 40 50 60 | 10 !0 !0 0 0 ) | 10 |
| Conventional Laboratory | Human (Sensititre Assisted) | Susceptibility Testing Result Reading (193-288) | | | | | | | nd Subculture (289-384): AST |
| RASP Laboratory | Robot (Micro) | | | | | | | | |
| | Robot (Prep) | | | | | | | | |
| | Human (Sensititre Assisted) | | | | | | | | |

| Day Fifteen | | Hour One | Hour Two | Hour Three | Hour Four | our Five | ur Six | r Seven | Eight |
|---|---|---|---|---|---|---|---|---|---|
| | | 10 20 30 40 50 60 | 10 20 30 40 50 60 | 10 20 30 40 50 60 | 10 20 30 40 50 60 | 10 20 30 40 50 60 | 10 20 30 40 50 60 | 10 !0 !0 0 0 ) | 10 |
| Conventional Laboratory | Human (Sensititre Assisted) | Antimicrobial Susceptibility Testing (289-384) | | | | | | | |
| RASP Laboratory | Robot (Micro) | | | | | | | | |
| | Robot (Prep) | | | | | | | | |
| | Human (Sensititre Assisted) | | | | | | | | |

| Day Sixteen | Hour One | Hour Two | Hour Three | Hour Four | Hour Five | Hour Six | Hour Seven | Hour Eight |
|---|---|---|---|---|---|---|---|---|

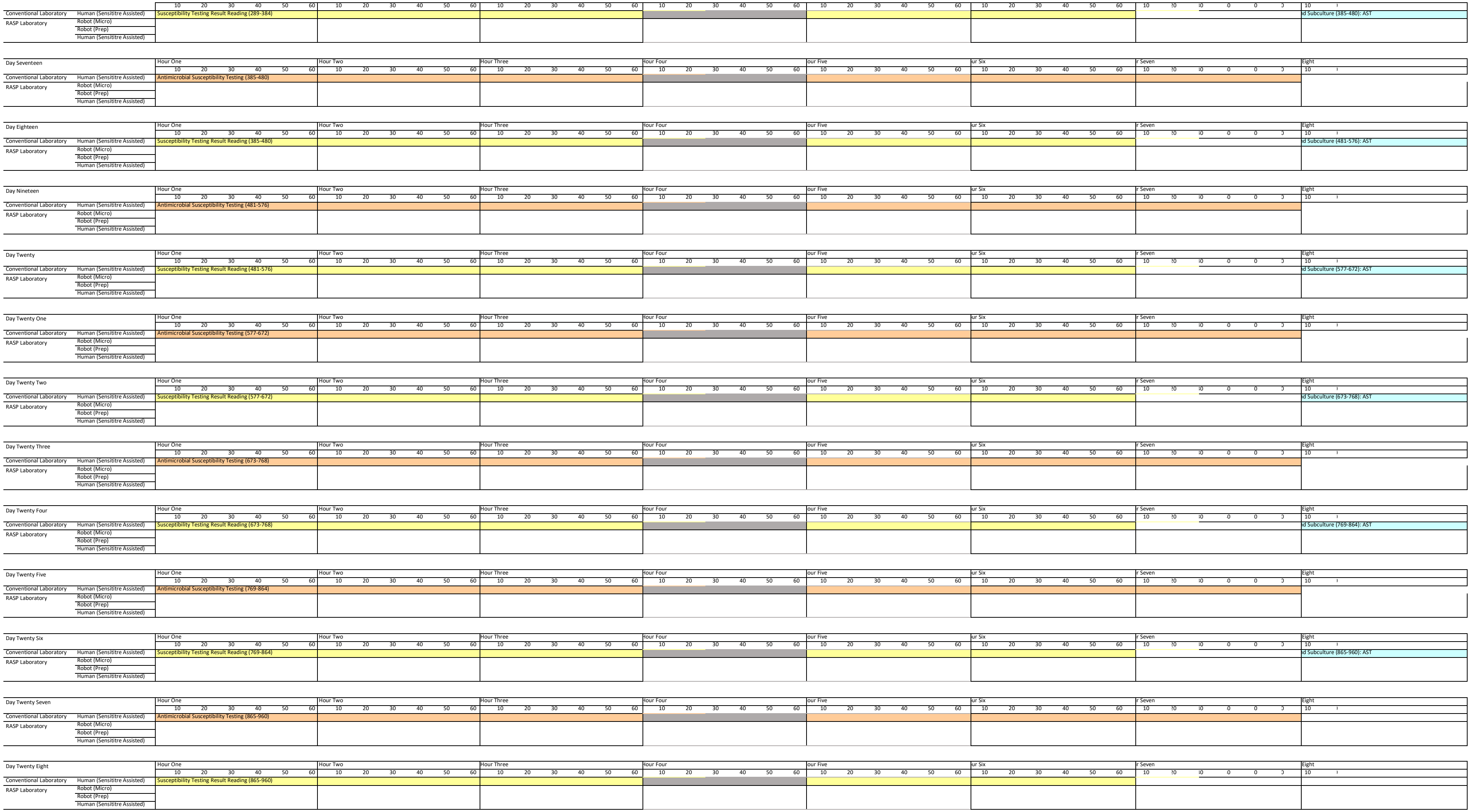


**Figure S3** | Time comparison simulation of a laboratory using the RASP robotic platform versus a conventional laboratory (each using only one technician) to process 960 isolates from homogenised faecal samples from isolation and identification through to antimicrobial susceptibility testing. The conventional laboratory was allowed the use of ‘modern’ laboratory implements such as Sensititre’s auto-inoculator and Vizion plate reader systems, pre-ordered ready-to-use drug plates, and a MALDI-TOF, while the RASP assisted laboratory had access to the RASP robotic platform and Tecan’s preparation robot (for drug plate preparation), a MALDI-TOF and a Sensititre Vizion plate reader system.
